# Dynamic Differences in Clinically relevant Pen β-Lactamases from *Burkholderia* spp

**DOI:** 10.1101/2024.11.02.621643

**Authors:** Jing Gu, Pratul K Agarwal, Robert A Bonomo, Shozeb Haider

## Abstract

Antimicrobial resistance (AMR) is a global threat, with *Burkholderia* species contributing significantly to difficult-to-treat infections. Ambler Class A Pen β-lactamases are produced by all *Burkholderia* spp., and their mutation or overproduction leads to the resistance of β-lactam antibiotics. This study investigates the dynamic differences among four Pen β-lactamases (PenA, PenI, PenL and PenP) using machine learning driven enhanced sampling molecular dynamics simulations, Markov State Models (MSMs), convolutional variational autoencoder-based deep learning (CVAE) and the BindSiteS-CNN model. Although sharing the same catalytic mechanisms, these enzymes exhibit distinct dynamic features due low sequence identity, resulting in different substrate profiles and catalytic turnover. The BindSiteS-CNN model further reveals local active site dynamics, offering insights into the Pen β-lactamase evolutionary adaptation. Our findings reported here identify critical mutations and proposes new hotspots affecting Pen β-lactamase flexibility and function, which can be used to fight emerging resistance in these enzymes.

## Introduction

Antimicrobial resistance (AMR) is a leading cause of death around the world (Antimicrobial Resistance, 2022). Key facts from WHO highlighted that AMR was the direct reason of 1.27 million global death in 2019 and contributed to 4.95 million deaths (Murray et al., 2022, Organization, 2023). Besides the high mortality rate, AMR was also estimated by the World Bank to result in the loss of US$ 1 to 3 trillion gross domestic product (GDP) annully by 2030 and cost US$ 1 trillion financial burden to the healthcare by 2050(Bank, 2017).

*Burkholderia* is a genus of Gram-negative bacteria that includes over 30 species of mammalian pathogens, some of which are clinically important to humans (Papp-Wallace et al., 2021, Becka et al., 2022). *Burkholderia cepacian* complex (BCC) and *Burkholderia gladioli* (B. *gladioli*) can infect individuals with cystic fibrosis (CF) and chronic granulomatous disease and cause difficult-to-treat chronic infections. *Burkholderia pseudomallei* (*B. pseudomallei*) is the etiologic agent of an often-fatal disease melioidosis, found in tropical and subtropical regions (Randall et al., 2016, Schweizer, 2012, Rholl et al., 2011). Outbreaks of BCC were reported in Hong Kong in 2020 (Cheng et al., 2024) and in the UK between 2023 to 2024 (Agency, 2024). In the US, as reported by CDC, *Burkholderia* related cases have been consistently reported between 2020 and 2024 (Vazquez Deida AA, 2024, Hudson MJ, 2022).

β-lactam antibiotics such as meropenem and ceftazidime are commonly recommended therapies for *Burkholderia* infections (Chirakul et al., 2018). However, the potency of these β-lactam antibiotics is declining because of the expression of bacterial β-lactamases that can destroy these drugs. All *Burkholderia* spp. can produce an Ambler class A Pen β-lactamase, whose overproduction and mutations might cause resistance to β-lactam antibiotics such as ceftazidime and amoxicillin-clavulanic acid (Schweizer, 2012, Papp-Wallace et al., 2021). PenA is a β-lactamase identified from the member of BCC named *Burkholderia multivorans.* It was found to be a carbapenemase most similar to KPC-2 (Trépanier et al., 1997, Papp-Wallace et al., 2013, Becka et al., 2022, Poirel et al., 2009, Becka et al., 2018). PenI produced by *B. pseudomallei* is also referred to as the soluble form of PenA and was shown to possess Extended-spectrum β-lactamase (ESBL) properties (Randall et al., 2016, Papp-Wallace et al., 2013, Papp-Wallace et al., 2016). PenL is a β-lactamase highly conserved in pathogenic *Burkholderia* spp. such as *B. pseudomallei*, *B. mallei*, and *B. cenocepacia* (called PenA in previous reports by H.S. Kim’s group (Yi et al., 2014, Hwang et al., 2014, Yi et al., 2012b, Yi et al., 2012a)) (Yi et al., 2016). PenL from *B. thailandensis* strain E264 confers resistance to amoxicillin (Yi et al., 2014). In *Burkholderia* spp., ceftazidime was also used for the selection of ceftazidime-resistant strains of *B. thailandensis* that express PenL (Papp-Wallace et al., 2017). PenP from *Bacillus licheniformis* is a narrow-spectrum class A β-lactamase that hydrolyses β-lactam antibiotics via the transient formation of an acyl-enzyme complex (Wong et al., 2011, Au et al., 2019).

Although all Pen β-lactamases share a common catalytic mechanism, their substrate profiles for a large number of clinically available antibiotics are very different (Papp-Wallace et al., 2013). By acquiring single or multiple mutations at certain strategic “hot-spots”, these enzymes experience rapid molecular evolution over a period of several years, leading to drastic expansion of their substrate profiles towards new generations of antibiotics (Wong et al., 2011). To reveal the indispensable structural and functional information related to such “hot-spots”, the sequence and structure differences between PenA (PDB ID: 3W4Q), PenI (PDB ID: 3W4P), PenL (PDB ID: 5GL9) and PenP (PDB ID: 6NIQ) were investigated, focusing on the dynamic differences between the four closely related Pen β-lactamases, using molecular simulations, Markov State Models (MSMs), and deep learning. More specifically, this work aims to differentiate and categorize conformations through convolutional variational autoencoder-based deep learning (CVAE) and the trained BindSiteS-CNN model, integrating binding site local similarity information into the representation of global protein dynamic properties. The results presented here highlight key residues and sites on Pen β-lactamases that have a potential impact on the flexibility and dynamic stability of the structure, eventually leading to the evolutionary functional differences.

## Results and Discussion

### Sequence and structure evolution

The Pen family of enzymes belong to class A β-lactamases according to the homology-based Ambler classification and have conserved secondary structures (Figure1A) (Papp-Wallace et al., 2021). However, there is considerable variation in their sequence identities, ranging from 51.33% to 89.81% (Figure1B). PenI and PenL share the highest sequence identity of 89.81%, which is consistent with being functionally similar as Extended-spectrum β-lactamase (ESBL). PenP and the other three Pen β-lactamases do not have very high sequence identity. The phylogenetic analysis revealed a possible evolutionary pattern amongst the Pen β-lactamases (Figure1C). The constructed phylogenetic tree posits PenP and PenA to be evolutionary close to each other, suggesting a common ancestor between them. In spite of all this, Pen β-lactamases still have conserved structural features like hydrophobic nodes and binding site residues (Figure1D).

### Structural and dynamic differences

The root means square fluctuation (RMSF) and deviation (RMSD) is often used as a measure to highlight flexibility in protein dynamics (Song et al., 2024). There are considerable dynamic structural differences observed from RMSF and RMSD analysis, observed between the four Pen β-lactamases (Figure 2, Figure S1). MSF comparison between the Pen enzymes was generated, using PenA as the reference structure (Figure 2A). The result has also been visualized using RMSF-coloured putty plots to highlight regions of differential flexibility (Figure 2B). The conventional RMSD fitting approach, which uses all Cα atoms, is ineffective in distinguishing between regions of high and low mobility in β-lactamases. To differentiate these regions, we used MDLovoFit analysis to perform RMSD fitting using a fraction (φ) of Cα atoms. Beyond this fraction, there is a sharp rise in the RMSD value for the remaining Cα atoms (Martínez, 2015). Such analysis can be effectively used to observe the least (blue) and the most mobile atoms (red) (Figure S1). Different RMSD and the corresponding fraction of aligned atoms (φ) were used for the four Pen β-lactamases (PenA: RMSD = 0.9 Å, φ = 0.79; PenI: RMSD = 1.0 Å, φ = 0.67; PenL: RMSD = 0.7 Å, φ = 0.72; PenP: RMSD = 0.55 Å, φ = 0.70). The analysis identifies several common dynamic motifs including the α3-α4 loop, Ω-loop (R164-D179), hinge region, α8-helix and the β9-α12 loop. Between the 4 enzymes, PenI is the most dynamic, while PenP is the most stable. The structure of PenA and PenI display similar dynamics with flexibility in the β9-α12 loop and the Ω-loop. While some flexibility is observed in α3-α4 loop in PenI, this is absent in PenA. To explore the correlated motions within the four Pen β-lactamases, we computed the dynamic cross-correlation maps (DCCMs) (Figure 2C, Figure S2) (Yu and Dalby, 2020). In these figures, the blue regions represent no to slightly negative correlations while light blue to green regions represent moderate positive correlations. Positive correlations represented with red colour imply residues moving in the same direction. There are more significant positive correlations identified in PenI (Figure S2), which suggest enhanced communication and cooperative motions across its structure while the least correlations in PenP reflect its stability observed.

**Figure 1.**
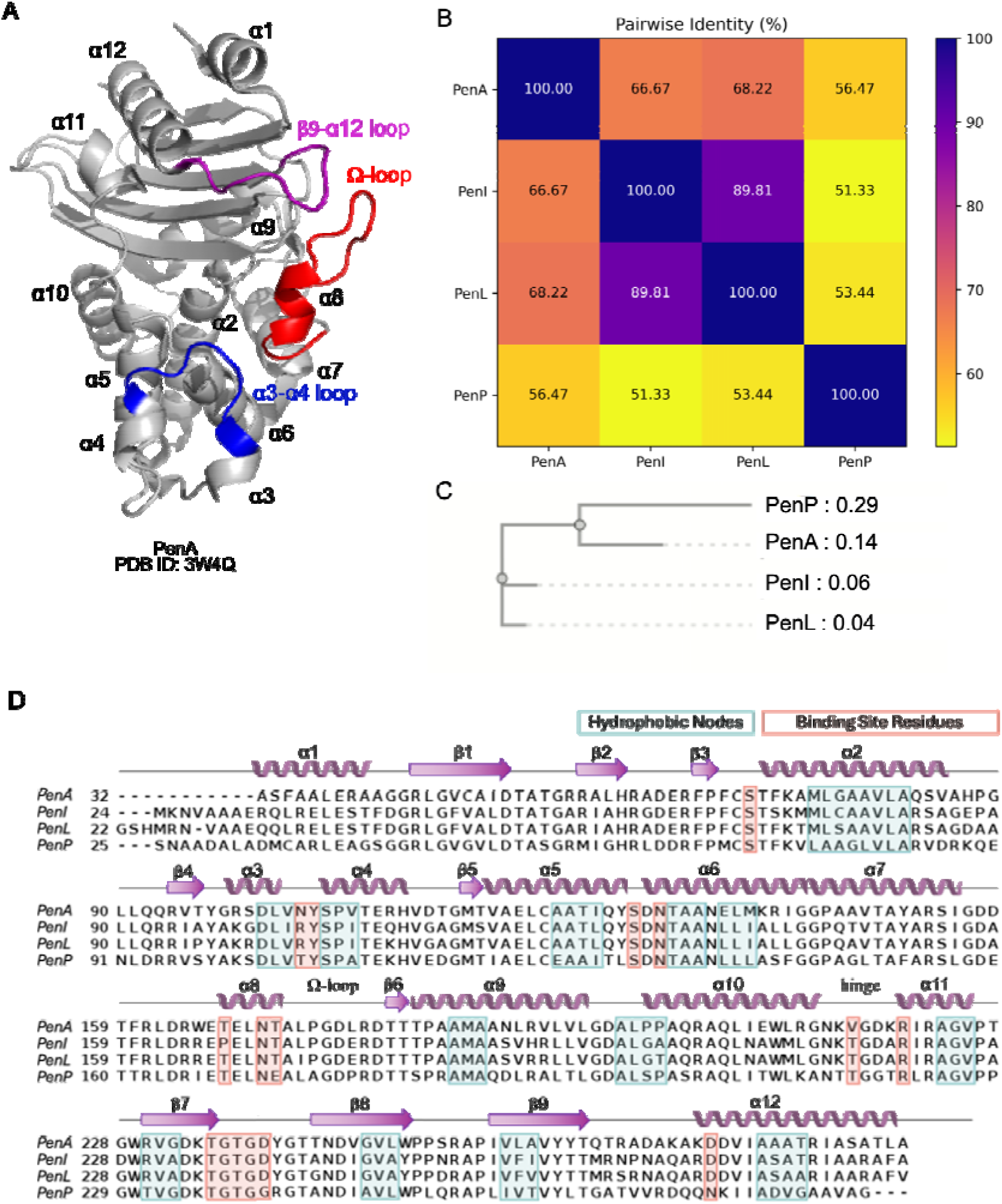
Structure and sequence of Pen β-lactamases. (A) Crystal structure of PenA β-lactamase (PDB ID: 3W4Q). (B) Pairwise sequence identity of the four Pen β-lactamases. (C) Phylogenetic tree of four Pen β-lactamases based on the multiple sequence alignment. (D) Multiple sequence alignments between PenA, PenI, PenL and PenP with secondary structure element annotations. The green boxes indicate the hydrophobic nodes, while orange boxes represent binding site residues.

**Figure 2.**
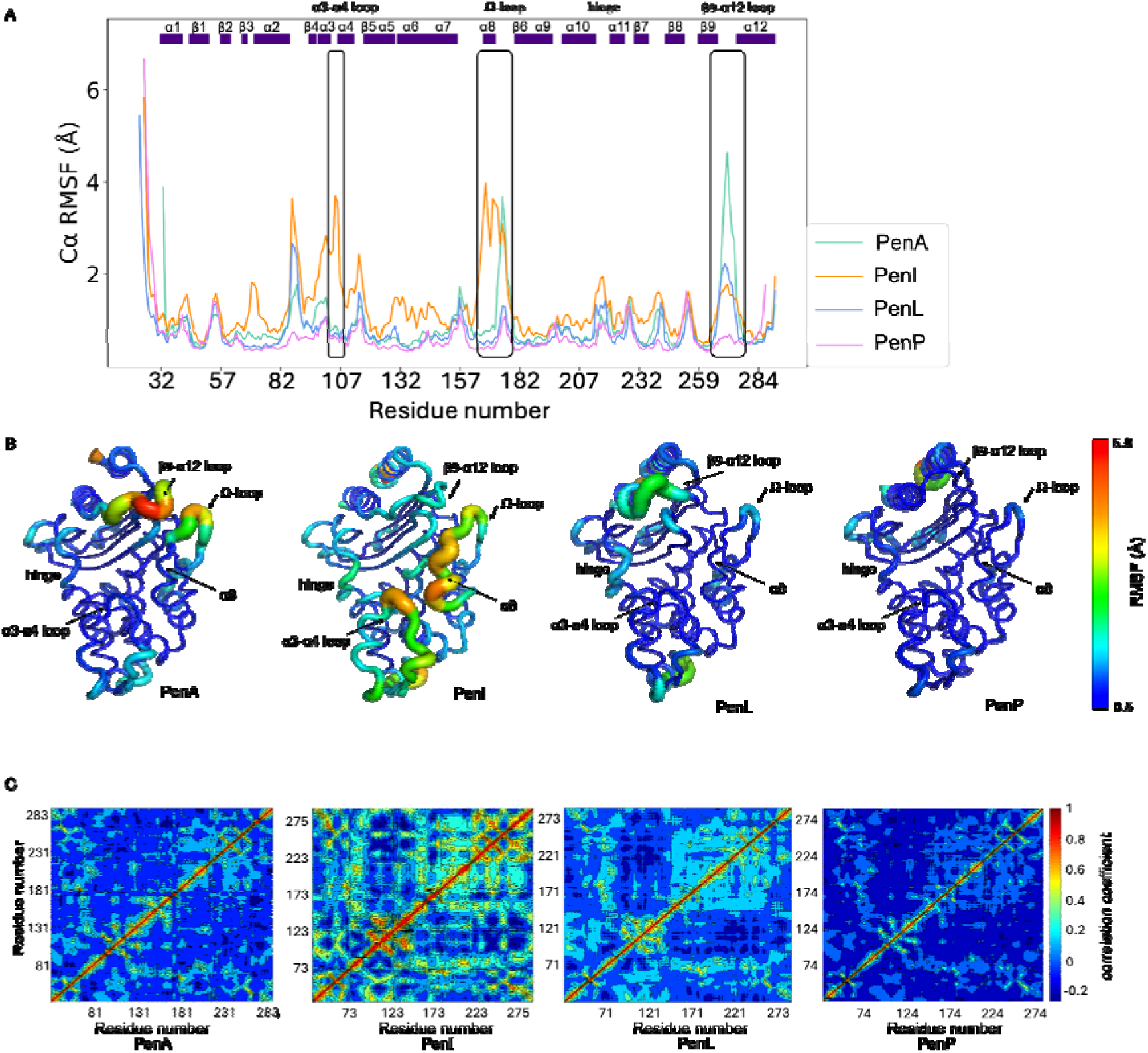
Conformational dynamics in Pen β-lactamases. (A) RMSF plots. (B) The putty structures of Pens. The putty structures are coloured based on the RMSF with the range clipped into 0.5 to 5.0 Å. (C) DCCMs.

To get a better understanding of the conformational differences, we further investigated the dynamics of these motifs in detail with MSMs and CVAE.

### Markov state models and Convolutional Variational Autoencoder

Four MSMs (Scherer et al., 2015) of the four Pen β-lactamases were successfully built using the backbone torsions of all residues and the χ1 angle from the residues of the hydrophobic nodes, α3-α4 loop, β9-α12 loop and the Ω-loop as input features (Figure S3-S6). The structural results were collected from each PCCA distribution state. For each state, 1000 frames were saved as the further input of CVAE-based deep learning analysis.

To further compare significant differences in structure conformations of Pen β-lactamases, the unsupervised CVAE-based deep learning approach was used (Bhowmik et al., 2018). Distance matrices of Cα of the hydrophobic node, α3-α4 loop, β9-α12 loop and the Ω-loop residues were calculated from the saved frames of all states of the four Pen β-lactamases and stacked together as the input data of CVAE. During the training stage of the CVAE, parallel experiments with the 3rd to the 30th latent dimension used were run. The model selected for decoding was the one with the lowest loss, which was with the 28th latent dimension.

A free energy landscape was generated based on the 2D PaCMAP representation of the CVAE latent space (Figure 3A). The four Pen β-lactamases show different dynamics and can be mainly clustered separately in different PaCMAP spaces while some conformations of PenA and PenI are clustered in the same space, which indicates that they share similar dynamics with these states. The ‘close/open’ conformations of Ω-loop and β9-α12 loop in PenA and the α8-helix and the α3-α4 loop in PenI are all observed by calculating the representative pairwise Cα distances: R65-G175 (Figure 3B), Y241-D271 (Figure 3C) in PenA, S70-P167 (Figure 3D), S70-Y105 (Figure 3E) in PenI. Since there are mutants with different amino acids in Pens, the calculated distances are of the pairs in the same position based on the alignment.

**Figure 3.**
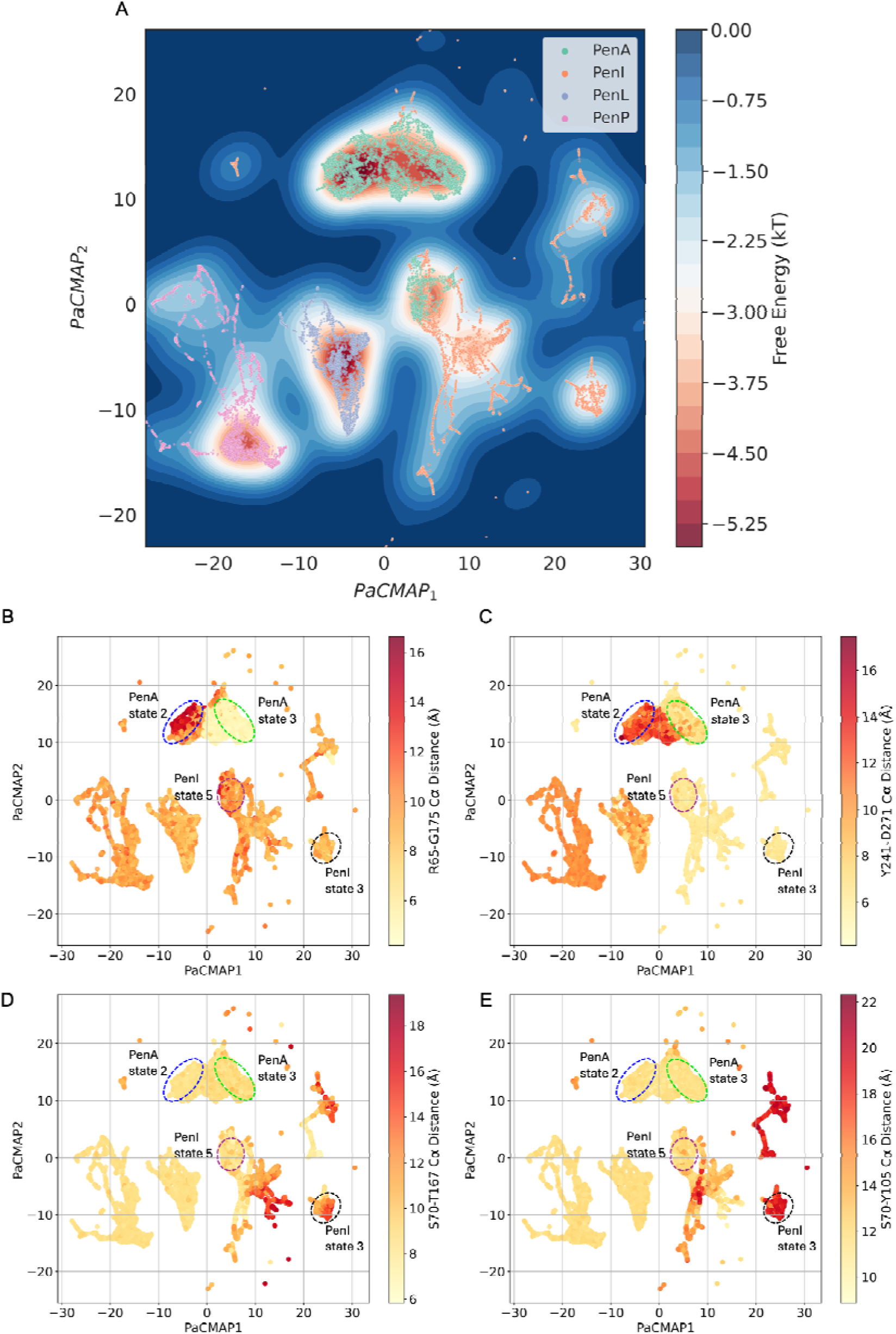
Convolutional variational autoencoder (CVAE)-based deep learning analysis. CVAE-learned features of high-dimensional data represented in 2D following PaCMAP treatment. (A) free energy landscape. (B) Cα distances of R65-G175 represent the conformation of the middle of Ω-loop. (C) Cα distances of Y241-D271 represent the conformation of β9-α12 loop. (D) Cα distances of S70-P167 represent the conformation of α8-helix. (E) Cα distances of S70-Y105 represent the conformation of the α3-α4 loop.

Referring to the distribution of states from the MSMs (Figure S7), state 2 of PenA represents the ‘open-open’ conformation of the middle of the Ω-loop and the β9-α12 loop, while state 3 of PenA represents the ‘close-close’ conformation of the Ω-loop and the β9-α12 loop (Figure 4A). State 3 of PenI represents the ‘open-open’ conformation of the α8-helix and the α3-α4 loop, while state 5 of PenI represents the ‘close-close’ conformation of the α8-helix and the α3-α4 loop (Figure 5A).

**Figure 4.**
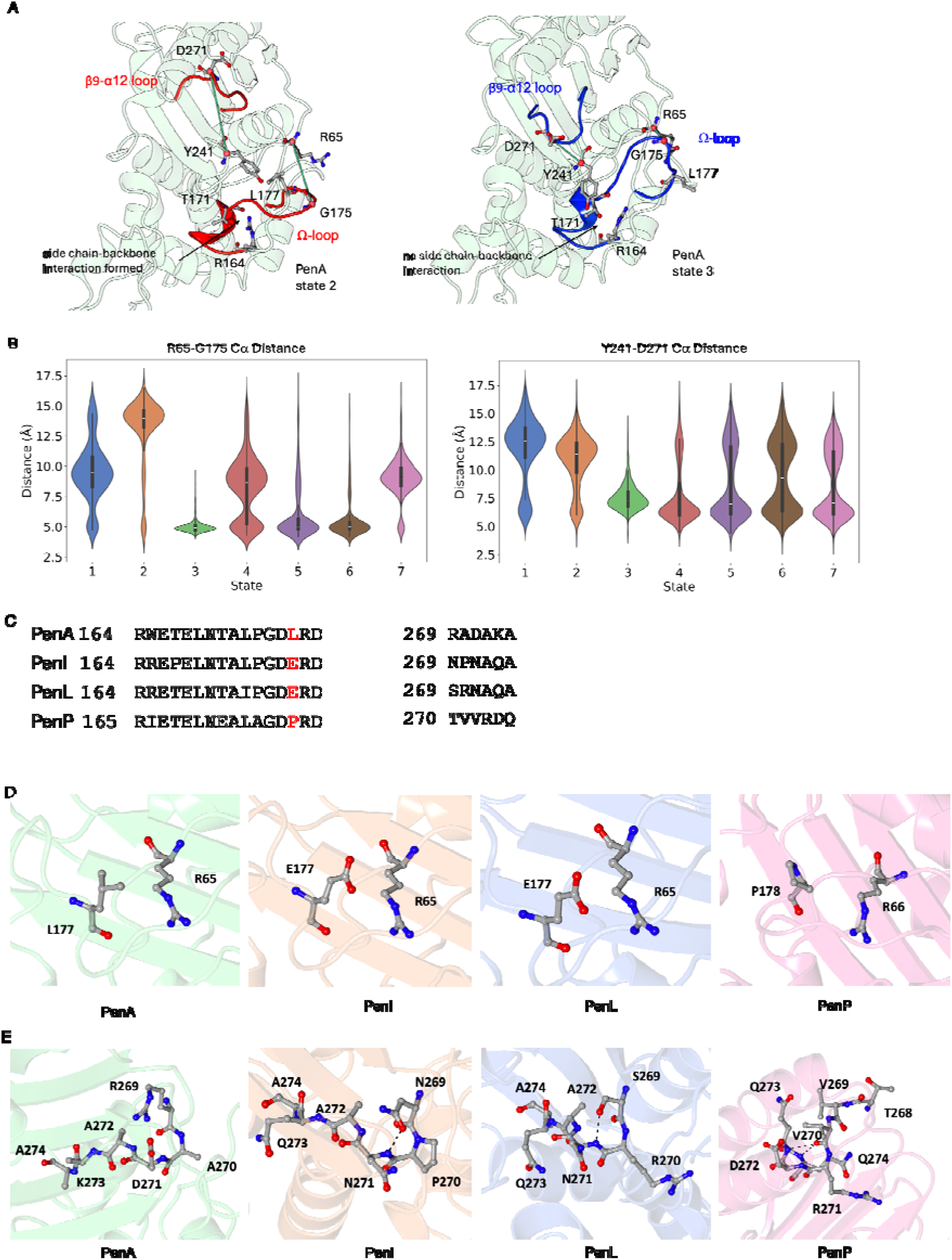
The β9-α12 loop and Ω-loop of Pen β-lactamases. (A) The open-open conformation of the central section of the Ω-loop and the β9-α12 loop from state 2 and the close-close conformation of them from state 3 of PenA. (B) Cα distance of R65-G175 and Y241-D271 in all PenA states. (C) Local sequence alignments of Pens. (D) L177 in PenA, E177 in PenI and PenL, P178 in PenP. (E) The β9-α12 loop of Pens.

**Figure 5.**
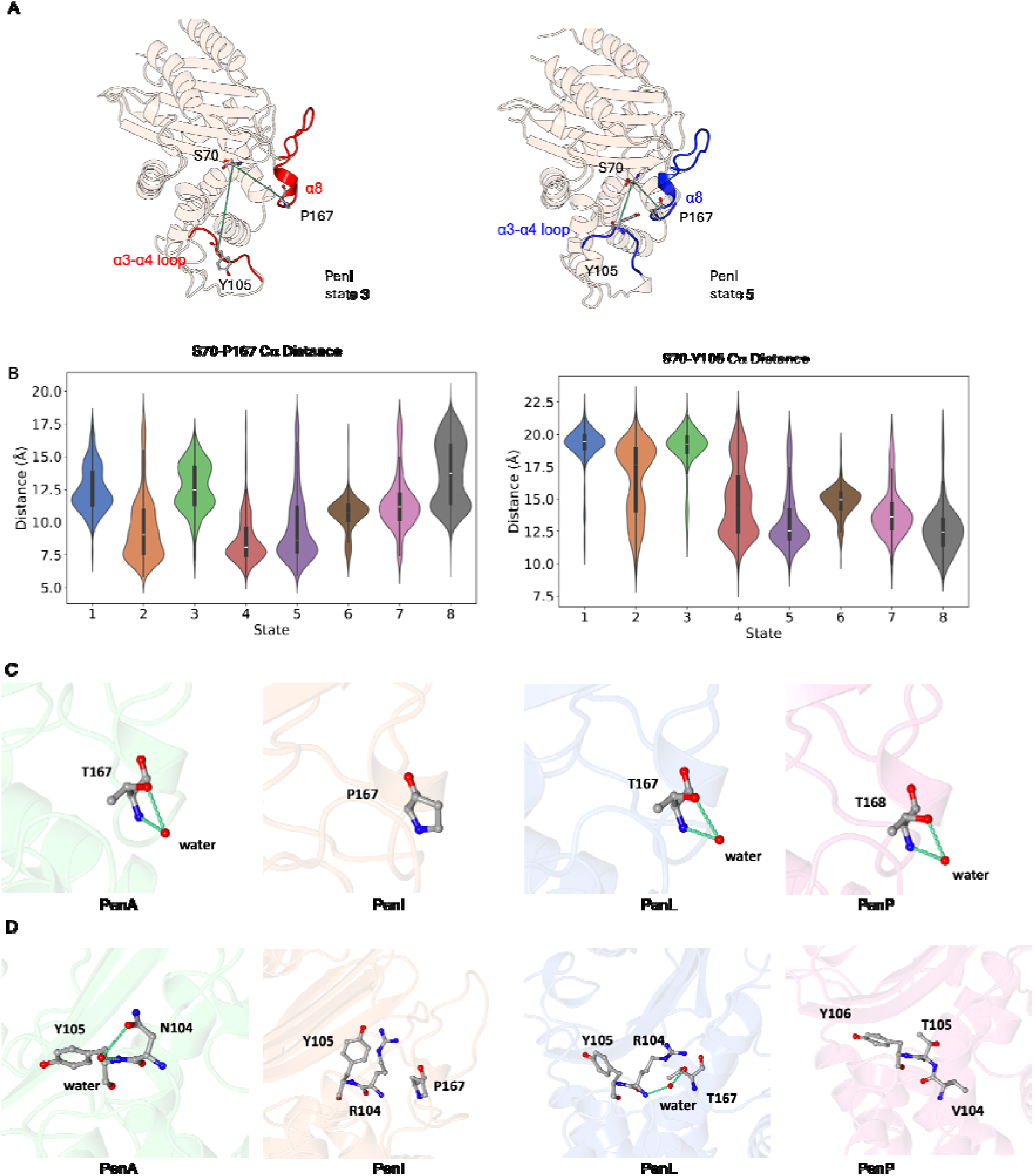
α8 helix and α3-α4 loop of Pen β-lactamases. (A) The open-open conformation of the central section of α8 helix and α3-α4 loop from state 3 and the close-close conformation of them from state 5 of PenI. (B) Cα distance of S70-P167 and S70-Y105 in all PenI states. (C) T167 in PenA, P167 in PenI, T167 in PenL, T168 in PenP. (D) The α3-α4 loop of Pens.

The central section of the Ω-loop (A172-R178) and the β9-α12 loop in PenA displays high flexibility with significant ‘open’ and ‘close’ conformations (Figure 4A). The calculation of the Cα distance of R65-G175 can represent the conformation of the Ω-loop since R65 is a stable site that can be used as the reference, while that of Y241-D271 can show the dynamic of the β9-α12 loop (Figure 4B). The central section of the Ω-loop in PenA and PenI is more dynamic than that of PenL and PenP (A173-R179) (Figure 2B). L177 in PenA might contribute to the dynamics in PenA (Figure 4C/D). The absence of E177-R65 interaction (Figure 4D) in PenA allows for a more flexible flip angle of the middle part of the Ω-loop. With the unique open state of PenA, side chain-backbone interaction between R164 and T171 can be formed and further stabilise the angle. Since the Ω-loop was reported to be very important to the binding of the substrate of β-lactamases, this extra flexibility might be one of the reason behind the different substrate preference of PenA.

Residue 269 positioned within the β9-α12 loop is poorly conserved between PenA (R269), PenI (N269), PenL (S269) and PenP (V270) enzymes (Figure 4C/E). In PenA, R269 forms an ion pair interaction with the side chain of D271 (Figure 4E). Similarly, the shorter side chains in PenI (N269) and PenL (S269) forms an H-bond interaction with the mainchain nitrogen of N271. In PenP, the structural arrangement of this loop is such that the side chain of T269 is unable to make any interactions. This structural arrangement instead allows the mainchain carbonyl oxygen of V270 to make two hydrogen bonds with the mainchain nitrogen atom of R272 and Q274 (Figure 4E). The residues to the α12-helix side of PenP can form more H-bond between the backbone of N270 and the backbone of D273 and Q274 (Figure 4E) that makes the α12-helix of PenP longer than that of PenA, PenI and PenL. This difference might be the reason for the higher stability of the β9-α12 loop in PenP.

The α8-helix is a short 3_10_ helix and just precedes the Ω-loop. The calculation of the Cα distance of S70-P167 can represent the conformation of the α8-helix since S70 is a stable site that can be used as the reference, while that of S70-Y105 can show the dynamic of the α3-α4 loop (Figure 5A/B). This helix in PenI is highly dynamic compared to that in PenA, PenL and PenP (Figure 2B). In PenI, a Proline is present at position 167, while this residue is a Threonine in PenA, PenL and PenP (Figure 5C). Proline is considered a potent breaker of alpha helices and beta sheets (Li et al., 1996). The presence of P167 in PenI significantly destabilises the α8-helix. The dynamic differences observed in the α3-α4 loop can also be explained by comparing PenI with other Pens (Figure 5D). In PenI, P167 is unable to make any interactions. However, T167 with a hydroxyl side chain can form interactions with the backbone of R104 via a stable bridging water molecule in PenL.

Residues E166 and N170 of the Ω-loop position a water molecule used during the acylation and deacylation of β-lactams (Figure 6A) (Au et al., 2019, Papp-Wallace et al., 2013). The presence of P167 in the α8-helix in PenI leads to a shift in the position N170 and subsequently results in an increase in the hydrogen bonding distance between E166 and N170. The hydrogen bond is formed only 16.2% of the simulation time (Figure 6B). Furthermore, the distance between N170 and the catalytic S70 also increases, thereby contributing to the overall increase in instability of the backbone in an indirect way (Figure 6C). The higher flexibility observed in PenI may explain the reason why PenI was also called the soluble form of PenA (Randall et al., 2016). This might result in the lower catalytic efficiency of PenI due to the weakening of interactions within the hydrogen bonding network associated with locating key water molecules within the binding site.

**Figure 6.**
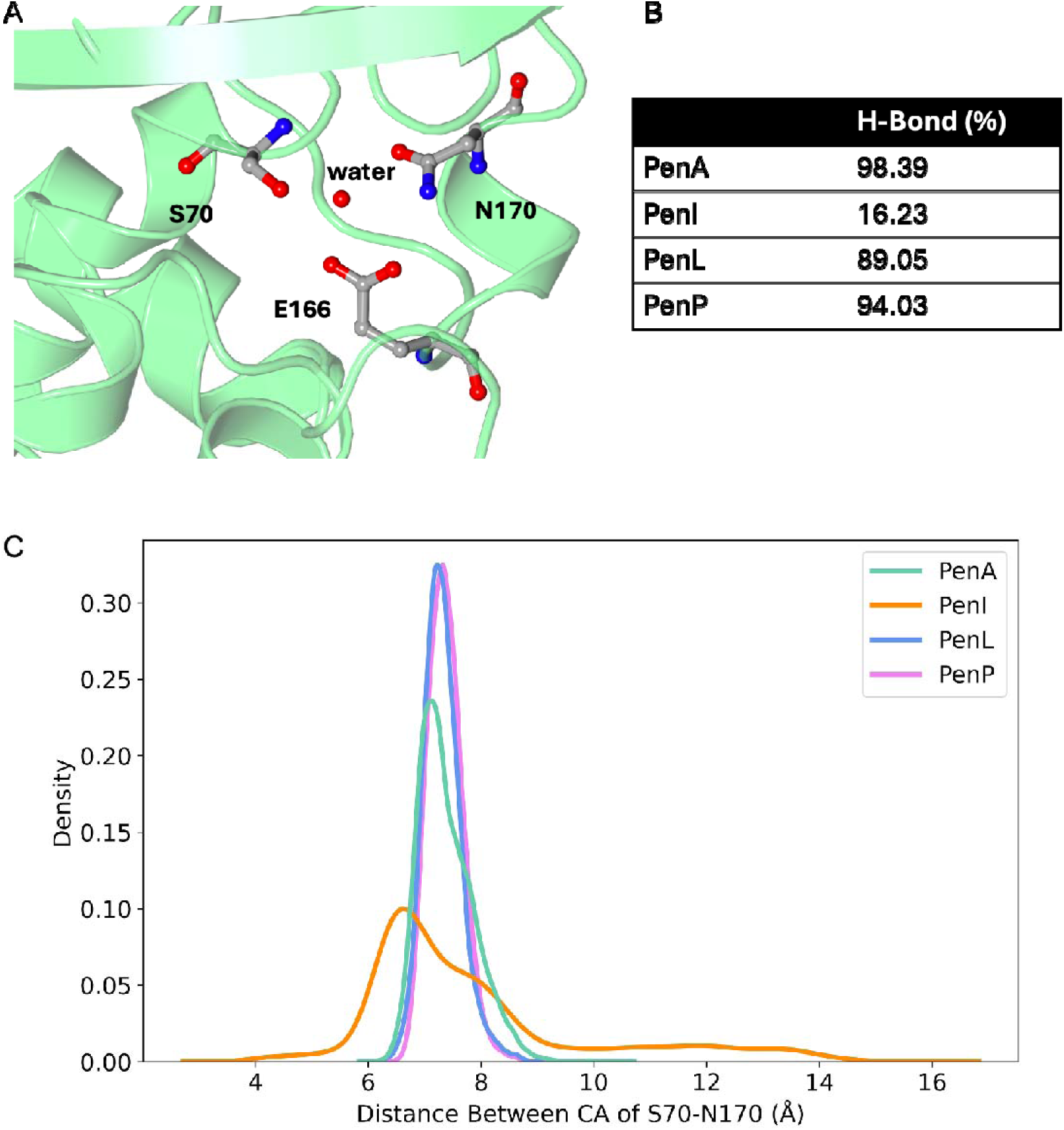
(A) Residues E166 and N170 of the Ω-loop position a water molecule used during acylation and deacylation of β-lactams. (B) The E166-N170 H-Bond ratio within the MD simulation. (C) The distance between the CA of S70 and N170.

The mutations that have been observed in the Pen family of β-lactamases, are also consistent with the evolutionary trend observed in other class A β-lactamases. The P167T substitution has been reported to cause ceftazidime resistance in CTX-M-23, another class A ESBL (Sturenburg, 2004). With T167, PenA has a one dilution higher level of ceftazidime resistance than PenI, which possesses P167 (Papp-Wallace et al., 2013). Furthermore, PenL containing T167 can be expressed in ceftazidime-resistant strains of *B. thailandensis* (Papp-Wallace et al., 2017). There is also evidence that PenP with T167 can bind with ceftazidime (Wong et al., 2011). The P167G substitution included in a triple mutant of another class A β-lactamase TEM-1, W165Y/E166Y/P167G, or the laboratory mutant P167S have also been reported to alter the conformation of the active site and results in ceftazidime hydrolysis (Stojanoski et al., 2015, Vakulenko and Golemi, 2002). These all suggest that position 167 is a hotspot in Pen β-lactamases that may alter conformational dynamics and impact function.

P174 was previously identified as one of the conserved residues within class A β-lactamases (Philippon et al., 2016). The emergence of A175 in the corresponding position in PenP indicates another possibility of substitution at this position. The flexibility of the Ω-loop is closely related to substrate selectivity in class A β-lactamases, and since PenP is a narrow spectrum β-lactamase, we speculate that P174 plays a key role in expanding the substrate range of the enzyme.

Residues at position 104 within Pen β-lactamases (N104 in PenA, R104 in PenI and PenL, T105 in PenP) have also been identified at analogous position in other representative class A β-lactamases as a hot spot. The point mutation D104E identified in SHV-1 β-lactamase can stabilize a key binding loop in the interface of SHV-1 and β-lactamase inhibitor protein (BLIP) by increasing the volume of the side chain of E104 (Reynolds et al., 2006). In TEM-1, the substitution of residue 104 has also been reported to produce extended-spectrum resistance and be related to the level of ceftazidime resistance (Vakulenko and Golemi, 2002).

Residue W105 was suggested to be important to the binding of β-lactams to KPC-2 and other class A β-lactamases (Papp Wallace et al., 2010). Previous studies of TEM-1 have also identified Y105 as a substitution hot spot (Reichmann et al., 2007). In L2 β-lactamases, the equivalent position is an H118Y substitution. This has been shown to affect the local environment within the active site (Zhu et al., 2024). Although Y105 is currently conserved in Pen β-lactamases, the orientation of Y105 has been reported to decrease the catalytic activity of PenI by obstructing the active site and preventing the β-lactam from binding (Papp-Wallace et al., 2013). Based on our hypothesis that class A β-lactamases might share a similar evolutionary trend, we can predict that position 105 could be a potential hot spot in Pen β-lactamases.

### BindSiteS-CNN based binding site comparison

BindSiteS-CNN is a Spherical Convolutional Neural Network (S-CNN) model trained to provide analyses of similarities in proteins based on their local physicochemical properties of their binding sites (Scott et al., 2022). It has been applied successfully to the study of L2 β-lactamases (Zhu et al., 2024). Our study is driven by the hypothesis that enzymes with similar structural features in the binding site will cluster together. The binding site residues were selected as highlighted in Figure 1D. Furthermore, the embeddings from the BindSiteS-CNN deep learning model ca be visualised in 2D with PaCMAP (Wang et al., 2021a). The input frames were taken from every 24th frame of the 298 trajectories in each system (PenA, PenI, PenL, and PenP).

Conformations of PenA, PenI, PenL and PenP extracted from the same energy basin with the most common area of overlap display high structural similarity (Figure7 i). This can also be considered as the most representative cluster of PenA. The following comparisons are referred to this set of relatively stable conformations. PenI is most flexible around the active site within the studied family of Pen β-lactamases. There are two significant clusters of PenI (Figure7 ii/iii. In the first cluster (CL1), the α3-α4 loop orients away from the active site, resulting in an enlarged active site (Figure10 ii/B). The calculated volume of the binding site within the simulation is 1074.34±204.16 Å^3^ for PenA, 1353.24±389.74 Å^3^ for PenI, 1329.58±170.27 Å^3^ for PenL, and 1079.47±131.60 Å^3^ for PenP (Figure 7C). In cluster 2 (CL2), the α8-helix moves away from the active site. This also results in an increase in the volume of the active site. In both clusters, the local features of the active site changes. (Figure7 ii/iii/B). The structure of the active site of PenP is stable and less flexible, resulting in a relatively smaller binding site. In PenP, the side chain of Y106 orients towards the hinge region rather than the Ω-loop (Figure10 iv). This might explain the narrow-spectrum activity of PenP. The exemplar conformation of PenL highlights the stable interactions between the side chain of T167 with the backbone of R104 via a bridging water molecule.

**Figure 7.**
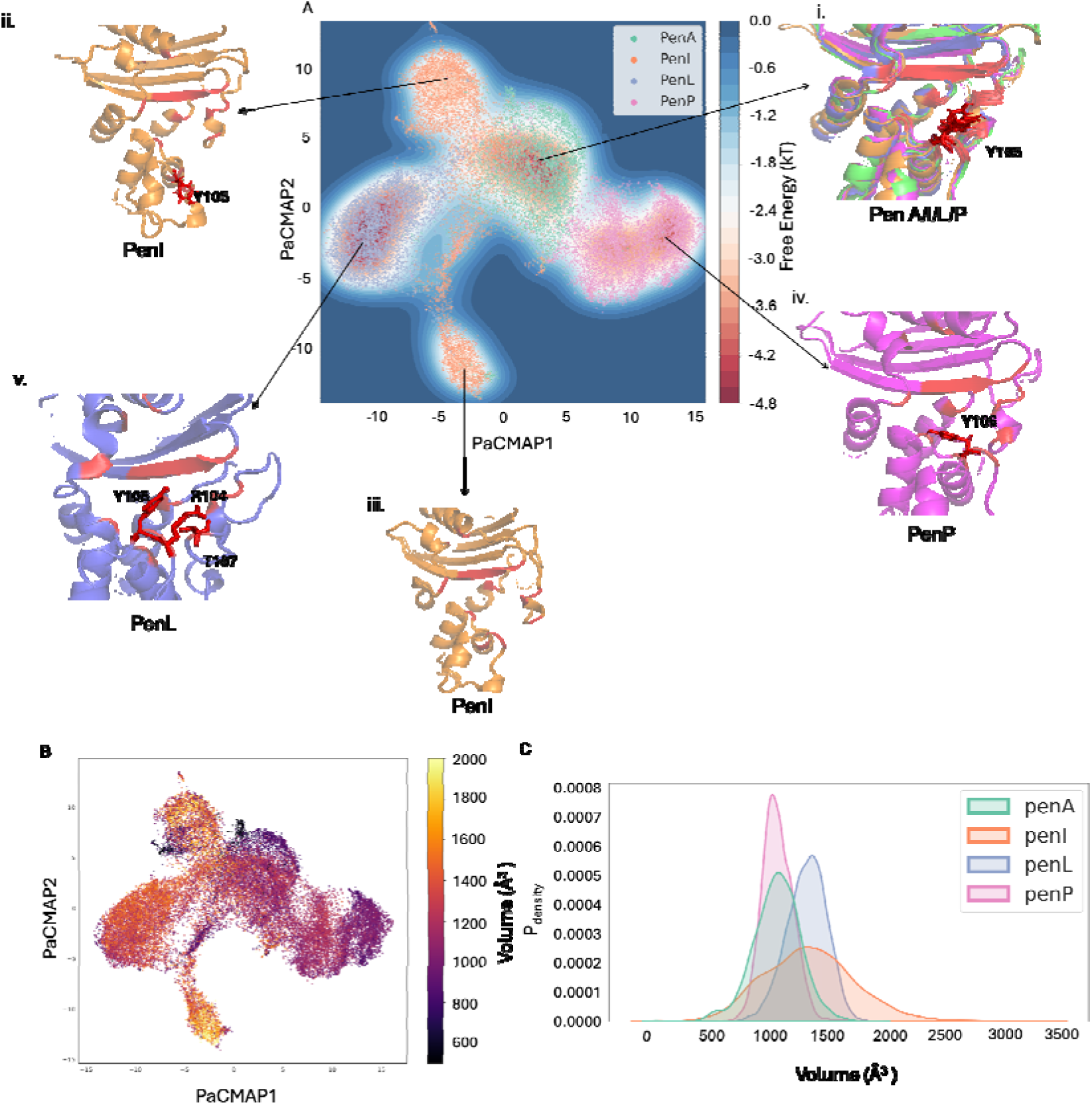
(A) BindSiteS-CNN-based high-dimensional embeddings represented in 2D with PaCMAP with free energy. (i) Conformations of PenA, PenI, PenL and PenP extracted from the same energy basin. (ii) The representative conformation of PenI with α3-α4 loop flipping away. (iii) The representative conformation of PenI with α8-helix flipping away from the active site. (iv) The representative conformation of PenP extracted. (v) Conformation was extracted from the main basin of PenL. (B) BindSiteS-CNN-based high-dimensional embeddings represented in 2D with PaCMAP colouring by binding pocket volume. (C) The calculated binding pocket volume of Pen β-lactamases

The PaCMAP of BindSiteS-CNN based embeddings highlight the differences between the local features of the active site in Pen β-lactamases. This analysis also reveals the local changes within the active site of Pen β-lactamases.

## Conclusions

To explore the dynamic differences between Pen β-Lactamases (PenA, PenI, PenL and PenP), this study employed adaptive sampling MD simulations, MSMs, CVAE and the BindSiteS-CNN deep learning model. Although the members are from the same Pen β-Lactamase family, we nevertheless captured several significant dynamic differences between them. The β9-α12 loop and Ω-loop are more dynamic in PenA due to the occurrence of L177 and R269. The α3-α4 loop, the Ω-loop, and the α8-helix are flexible in PenI. The emergence of P167 in PenI significantly improves its flexibility and solubility by disrupting the secondary structure of α8-helix. This alters the interactions made with the conserved water molecule required in the deacylation step of catalysis. The high level of structural stability of PenP is also consistent with its narrow spectrum activity. The application of the BindSiteS-CNN model to compare the local active site dynamics of Pens revealed the difference between all four simulated systems. The potential hot spots highlighted in this study may serve as a guide for further understanding of the biological functions and the evolutionary relationship of Pen β-Lactamases.

## Materials and methods

### Structural Models

The crystal structures of PenA (PDB ID 3W4Q) (Papp-Wallace et al., 2013), PenI (PDB ID 3W4P) (Papp-Wallace et al., 2013), PenL (PDB ID 5GL9) (Yi et al., 2016), and PenP (PDB ID 6NIQ) were downloaded from the Protein Data Bank.

### Sequence Alignment and Phylogenetic Tree Generation

Amino acid sequences of PenA, PenI, PenL and PenP were derived from the structural files through systematic parsing of the respective PDB structure files through pdb2fasta. A multiple sequence alignment was conducted utilizing Clustal Omega employing its default parameters (Madeira et al., 2022). The resulting alignment was subsequently visualized and interpreted with Jalview version 2.11.3.3 (Waterhouse et al., 2009) to furnish a graphical depiction of sequence congruities for improved comprehension.

### Systems preparation and Adaptive Sampling MD simulations

Molecular dynamics simulations of PenA, PenI, PenL and PenP were conducted with the following protocol. The initial system preparation utilized the PlayMolecule ProteinPrepare Web Application at pH 7.4 (Martinez-Rosell et al., 2017). All heteroatoms were excised from the PDB files. ProteinPrepare autonomously executed pKa calculations and optimized hydrogen bonds, while simultaneously assigning charges and protonating the PDB file within the high-throughput molecular dynamics (HTMD) framework (Doerr et al., 2016). Charge assignments were based on the local environment of the protonated structure, optimizing its hydrogen-bonding network. Utilizing tleap from the Amber MD package (Case et al., 2023), input files containing detailed information about atoms, bonds, angles, dihedrals, and initial atom positions were generated. The Amberff14SB force field (Maier et al., 2015) was applied, and each system was solvated in TIP3P water model (Mark and Nilsson, 2001) within a cubic box, maintaining a minimum 10 Å distance from the nearest solute atom, and neutralized with Na^+^ and Cl^-^ ions. Prepared systems were initially minimized through 1,000 iterations of steepest descent and subsequently equilibrated for 5 ns under NPT conditions at 1 atm, employing the ACEMD engine within the HTMD framework (Doerr et al., 2016, Harvey et al., 2009). The temperature was steadily increased to 300 K with a time step of 4 fs, using rigid bonds, a 9 Å cutoff, and particle mesh Ewald summations (Cerutti et al., 2009, Wells and Chaffee, 2015) for long-range electrostatics. During equilibration, the protein backbone and ligand atoms were restrained with a spring constant of 1 kcal mol^−1^Å^−2^, while the Berendsen barostat (Feenstra et al., 1999) controlled pressure and velocities were based on the Boltzmann distribution. The production phase, conducted in the NVT ensemble, used a Langevin thermostat with 0.1 ps damping and a hydrogen mass repartitioning scheme, allowing a 4fs time step and recording trajectory frames every 0.1 ns, all culminating in a final, unrestrained production step to observe natural system dynamics. The simulations were run until a minimum of 298 trajectories were obtained, with each trajectory counting 600 frames and ampling a cumulative 17.88 μs for each system.

### Markov State Models

Pyemma v2.5.12 was used to build the MSMs (Scherer et al., 2015). Backbone dihedral angles (φ and ψ) of all residues and the 1 angle from the residues of the hydrophobic nodes, α3-α4 loop, β9-α12 loop and the Ω-loop of Pens were selected as the input features. The featurised trajectories were projected onto selected independent components (ICs) using TICA. The produced projections can show the maximal autocorrelation for a given lag time. The chosen ICs were then clustered into selected clusters using k-means. In this way, each IC was assigned to the nearest cluster centre. The lag time was chosen to build the MSM with metastable states according to the implied timescales (ITS) plot. After passing the Chapman-Kolmogorov test within confidence intervals, the MSM was defined as good. This indicates the model highly agrees with the input data, and it is statistically significant for use. Bayesian MSM was used to build the final model in the system. The net flux pathways between microstates, starting from state 1, were calculated using Transition Path Theory (TPT) function. The pathways all originate from state 1, as it shows the lowest stationary probability (the highest free energy) in the system. This is why state 1 is a reasonable starting point to illustrate all the relevant kinetic transitions through the full FE landscape. The structural results were selected from each PCCA distribution.

### Deep conformational clustering using CVAE

The utilisation of CVAE was implemented in a systematic manner to reveal the dynamic difference of Pen β-lactamases. Data derived from each protein system such as the pairwise distances were computed using MDAnalysis (Michaud-Agrawal et al., 2011) and MDTraj (McGibbon et al., 2015). Detailed analyses involved the formulation of pairwise distance maps extracted from all states of the MSM of each system. The focus was directed towards hydrophobic nodes, α3-α4 loop, β9-α12 loop and the Ω-loop residues featuring the constraint of Cα atom distances ≤8 Å, which were recorded as non-zero values in a specified three-dimensional matrix. The cumulative data, represented as 74 × 74 distance matrices, were consolidated into a unified 3D matrix for every system, accompanied by a label file containing pertinent metadata.

The CVAE’s encoding section was structured with an 80:20 validation ratio and underwent training across 100 epochs. Dimensions spanning from 3 to 30 were explored, eventually settling on the 28th dimension, which exhibited the minimal loss, for the model’s architecture. Continuous oversight was maintained for potential overfitting. For the decoding component, matrices and label files derived from Pens were used to assess the model’s performance and to discern the clustering patterns of the conformations inherent to these systems.

The PaCMAP algorithm was then employed to reduce the dimensionality of the decoded embedding into two dimensions, thereby simplifying visualisation. This, when combined with the free energy landscape, proved instrumental in isolating distinctive conformations from energetically favourable regions.

### BindSiteS-CNN based binding site comparison

BindSiteS-CNN was employed to capture the differences between the active site local features of the four systems: PenA, PenI, PenL and PenP. The methodology encompassed binding pocket surface preparation and BindSiteS-CNN model processing and was adopted from Scott et al. (Scott et al., 2022). The samples were taken every 24th frame of the 298 trajectories in each system.

In the binding pocket surface preparation phase, the binding pocket surface of each frame was generated with side chain atoms of the binding site residues as the filtering reference. The computed pocket surface meshes with vertices enriched with physicochemical information describing the hydrophobicity, electrostatic potential and interaction-based classification of surface-exposed atoms lining the pocket were saved as PLY files and integrated as part of an in-house β-lactamases active pocket database.

During the BindSiteS-CNN model processing stage, the prepared 3D pocket mesh objects were fed into the trained BindSiteS-CNN model as input data. The embeddings of all input frames from the BindSiteS-CNN model as output data were saved out with labels into one pkl file. PaCMAP (Wang et al., 2021b) has been used to visualize their distribution in the descriptor space by downgrading high-dimensional embeddings to 2D, those dots represent similar binding sites would cluster together.

### Structural Analysis

The trajectories of the molecular simulations were meticulously aligned to their corresponding structures with MDAnalysis (Michaud-Agrawal et al., 2011) and MDTraj (McGibbon et al., 2015). The stride of frames within these trajectories was retrieved using the identical set of tools. To elucidate the general dynamics features inherent in the trajectories, calculations were performed again leveraging the functions within the MDTraj and MDAnalysis packages and MDLovofit (Martínez, 2015). For a more visual and intuitive understanding, the trajectories were loaded into the PyMOL Molecular Graphics System. This tool also facilitated the superimposing of structures and enabled a comprehensive conformational comparison. After delineating the spatial variations between distinct conformations, visual representations were generated via the Protein Imager (Tomasello et al., 2020). Additionally, the Matplotlib package (Hunter, 2007) in Python was employed for all statistical and graphical representations, including plots and figures, to present the data in a comprehensive and interpretable manner.

