## Supplementary Information for "Dynamic Differences in Clinically relevant Pen β-Lactamases from *Burkholderia* spp"

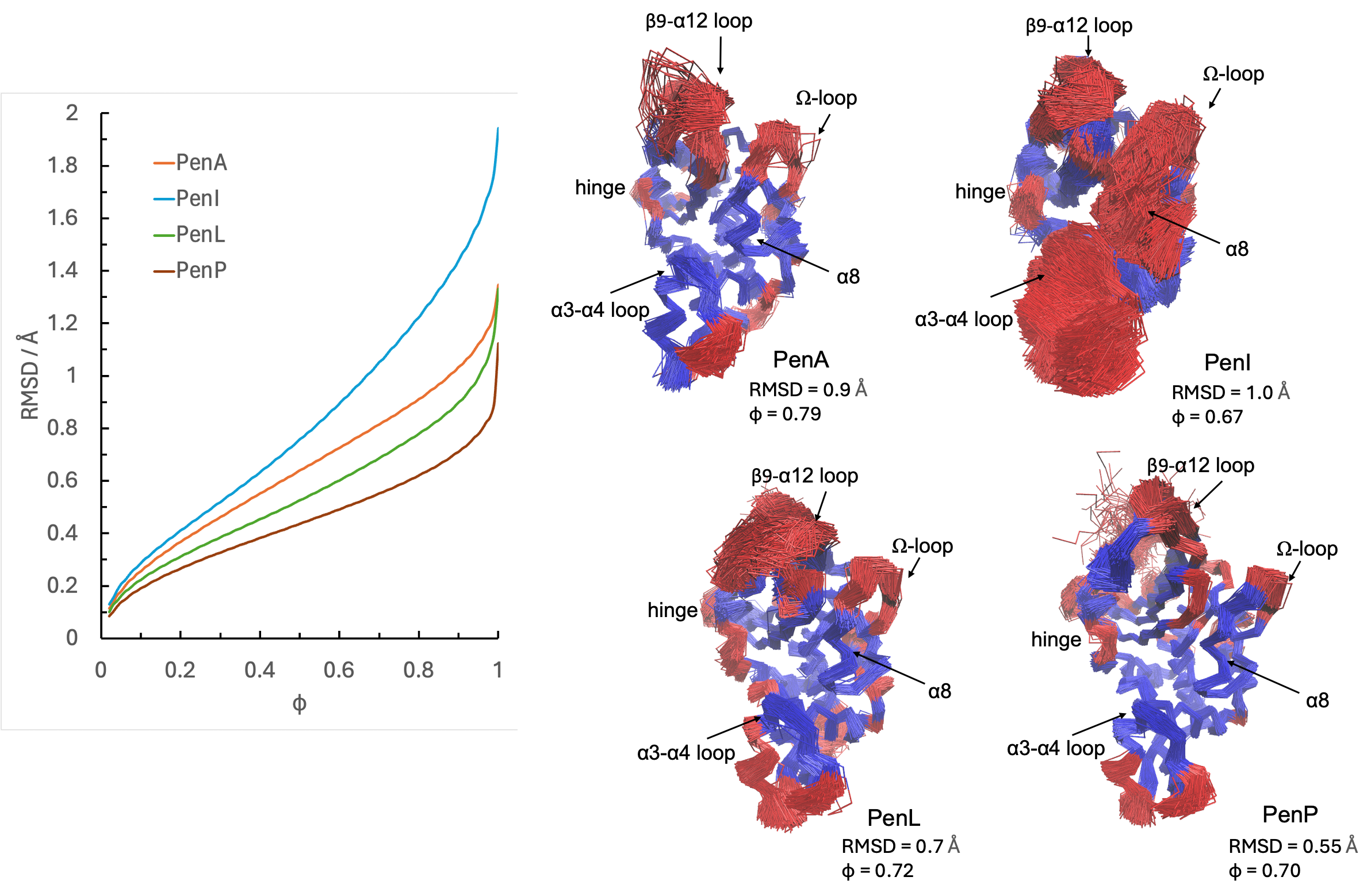

**Figure S1** Core Cα RMSD superimposition from Pen simulations. The structural alignment was calculated from all trajectories with stride 24 and rendered to illustrate 745 uniformly separated frames. The least mobile Cα atoms are coloured blue and the most mobile atoms (red) provide the structural basis for the differential RMSDs.

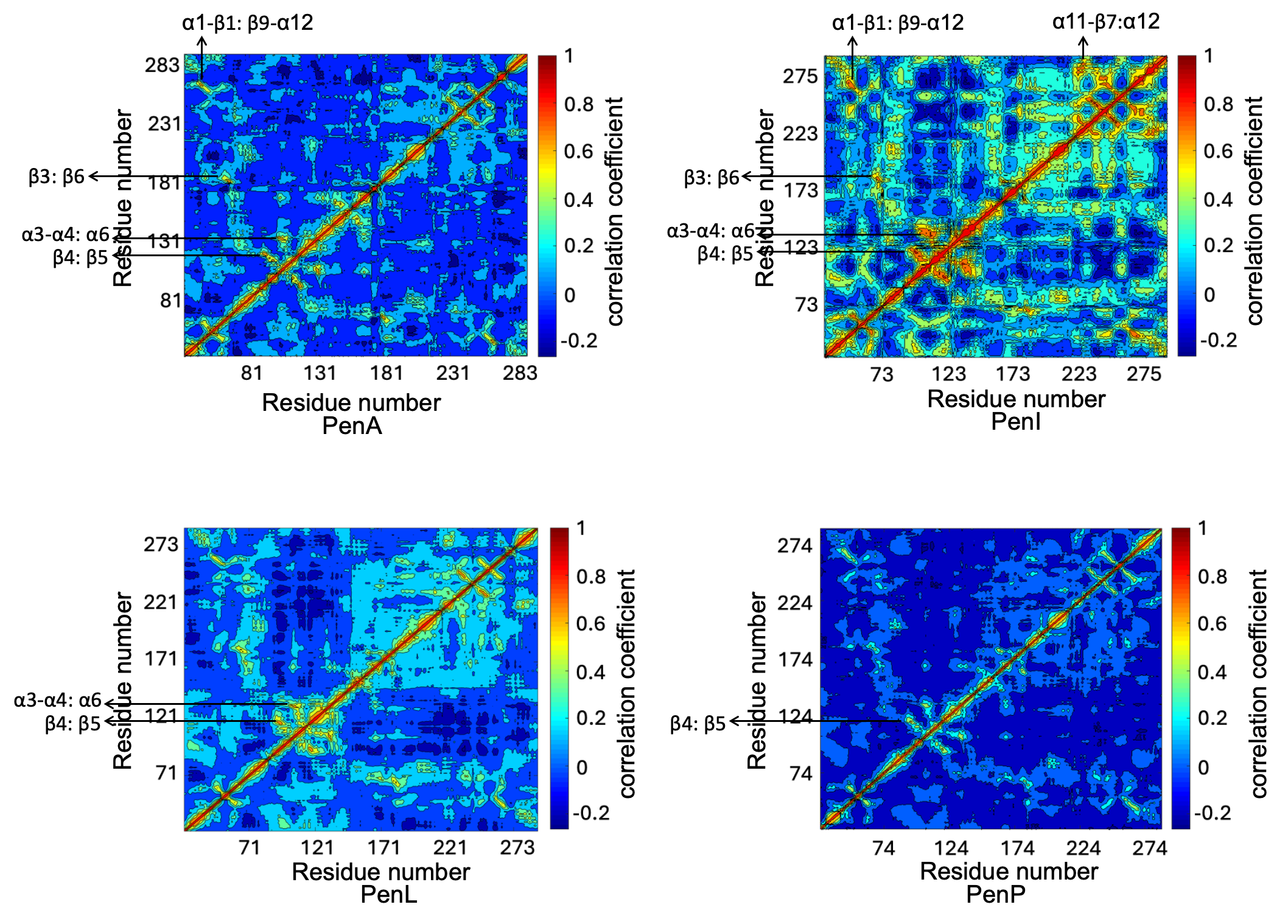

**Figure S2** Dynamic cross-correlation map (DCCM) computed from Pen simulations. The regions showing significant positive correlations are highlighted.

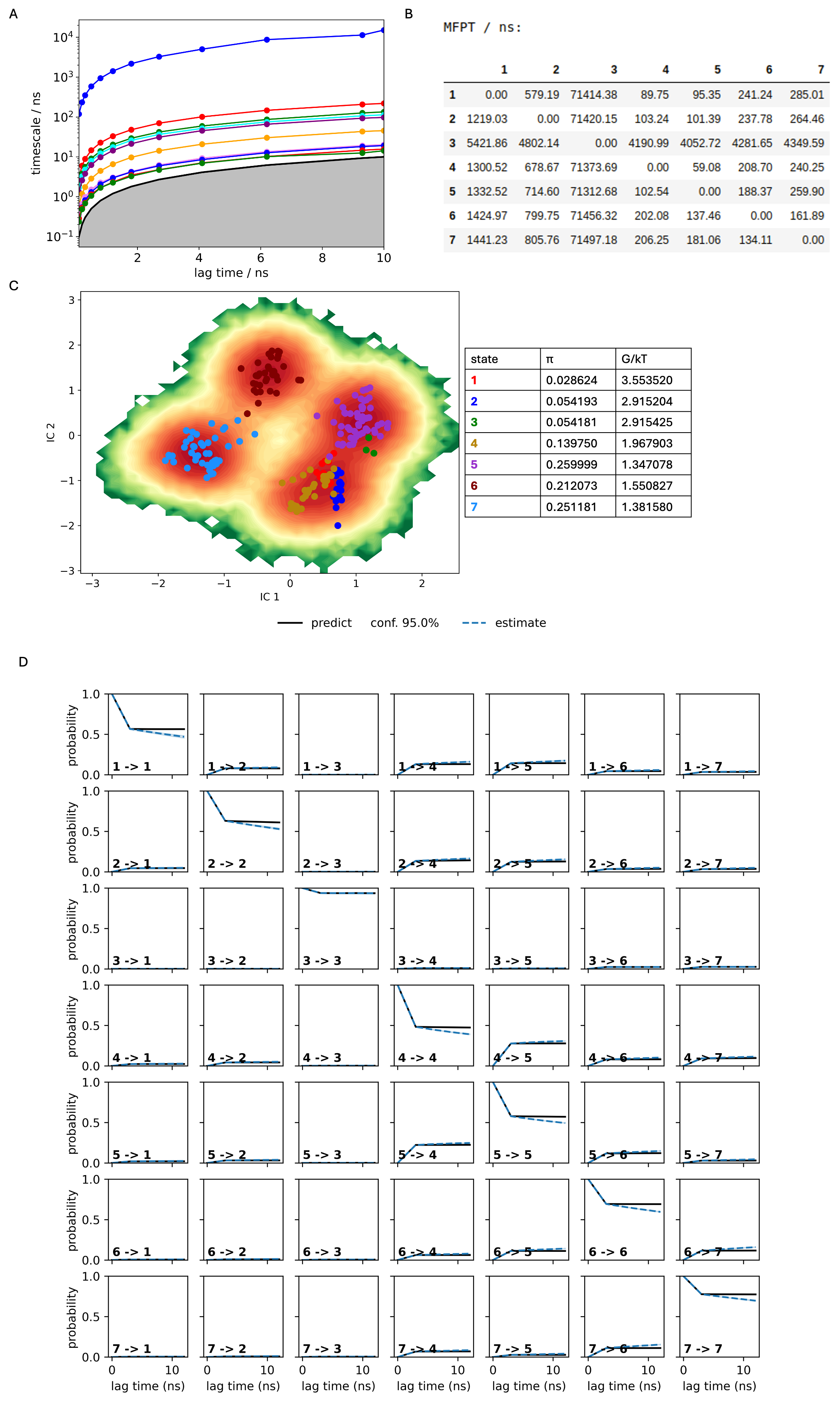

**Figure S3** **PenA Markov State Model.** A. Implied timescales (ITS) plot. B. Mean first passage times between metastable states per ns. C. The distribution of cluster centres highlighting the presence of the metastable states. D. Chapman-Kolmogorov (CK) test plots

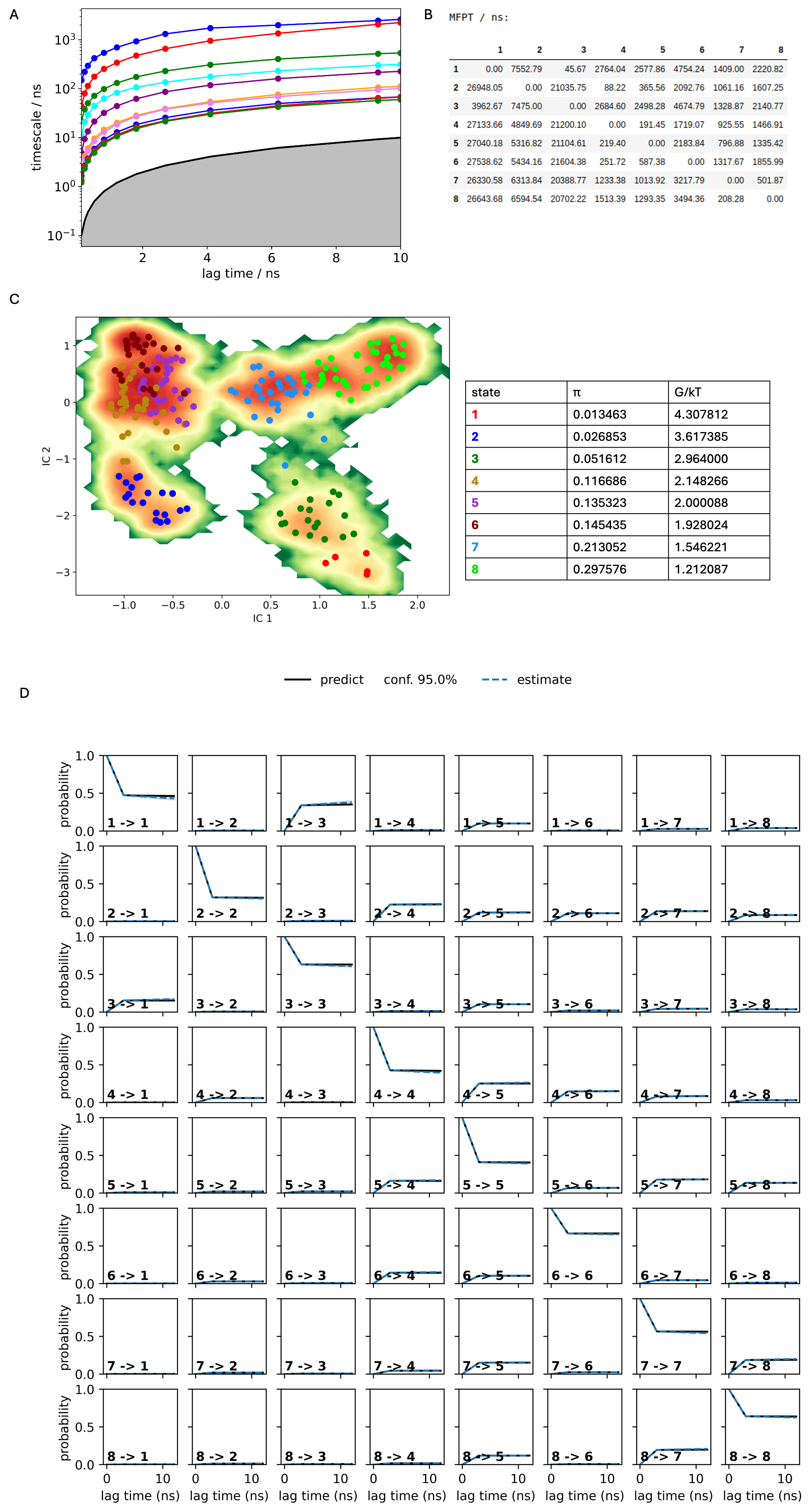

**Figure S4** **PenI Markov State Model.** A. Implied timescales (ITS) plot. B. Mean first passage times between metastable states per ns. C. The distribution of cluster centres highlighting the presence of the metastable states. D. Chapman-Kolmogorov (CK) test plots

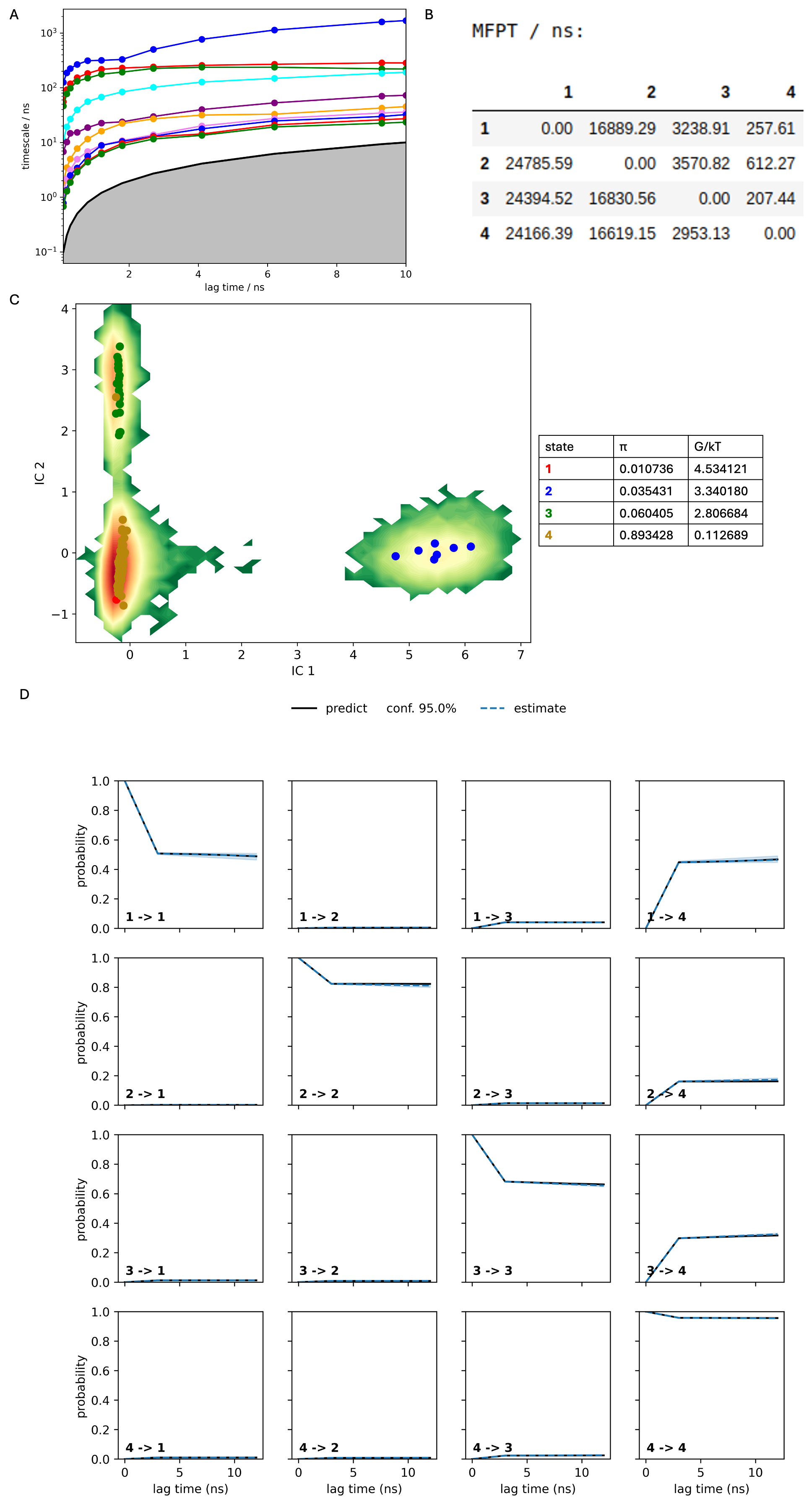

**Figure S5 PenL Markov State Model.** A. Implied timescales (ITS) plot. B. Mean first passage times between metastable states per ns. C. The distribution of cluster centres highlighting the presence of the metastable states. D. Chapman-Kolmogorov (CK) test plots

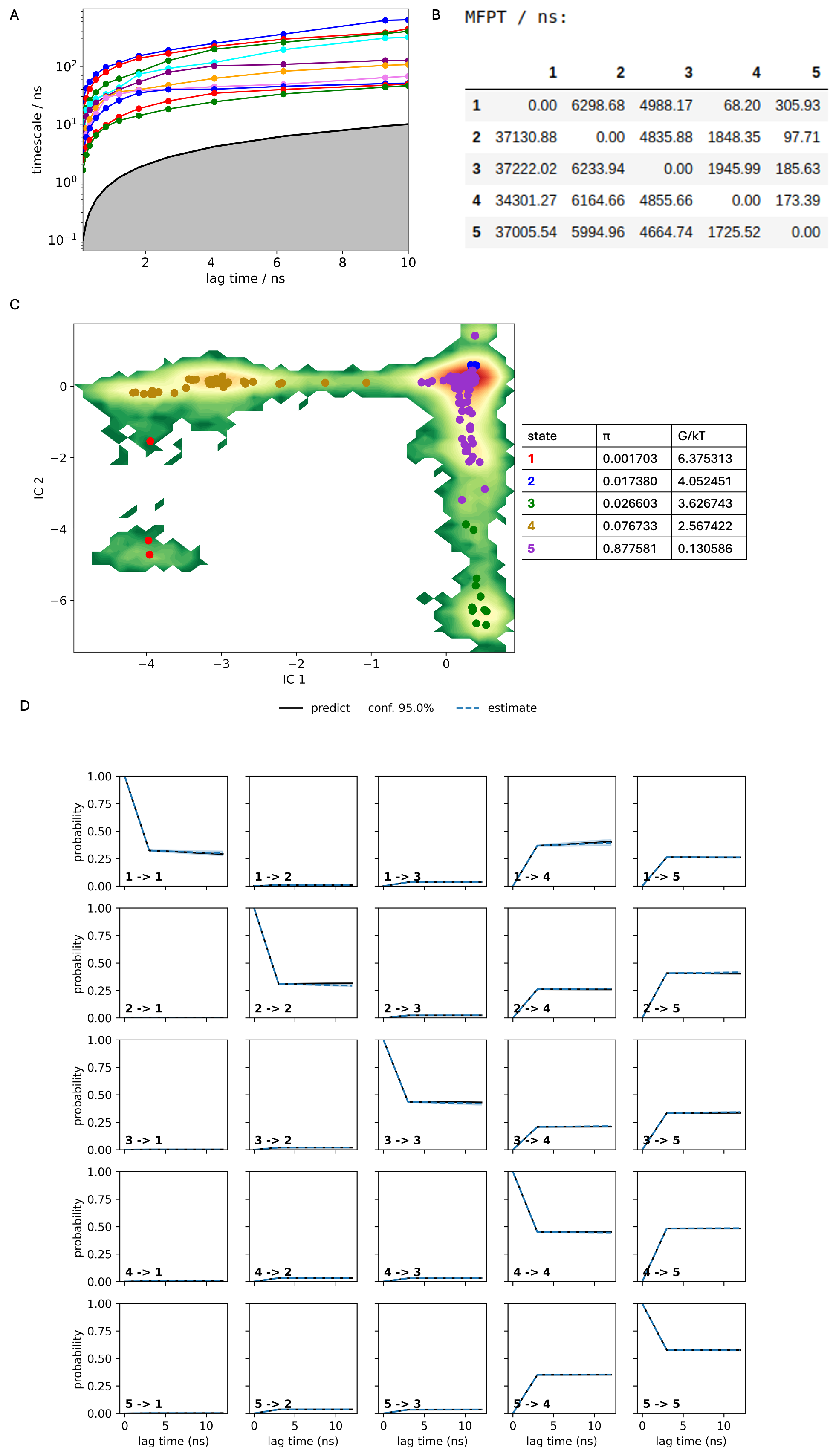

**Figure S6 PenP Markov State Model.** A. Implied timescales (ITS) plot. B. Mean first passage times between metastable states per ns. C. The distribution of cluster centres highlighting the presence of the metastable states. D. Chapman-Kolmogorov (CK) test plots

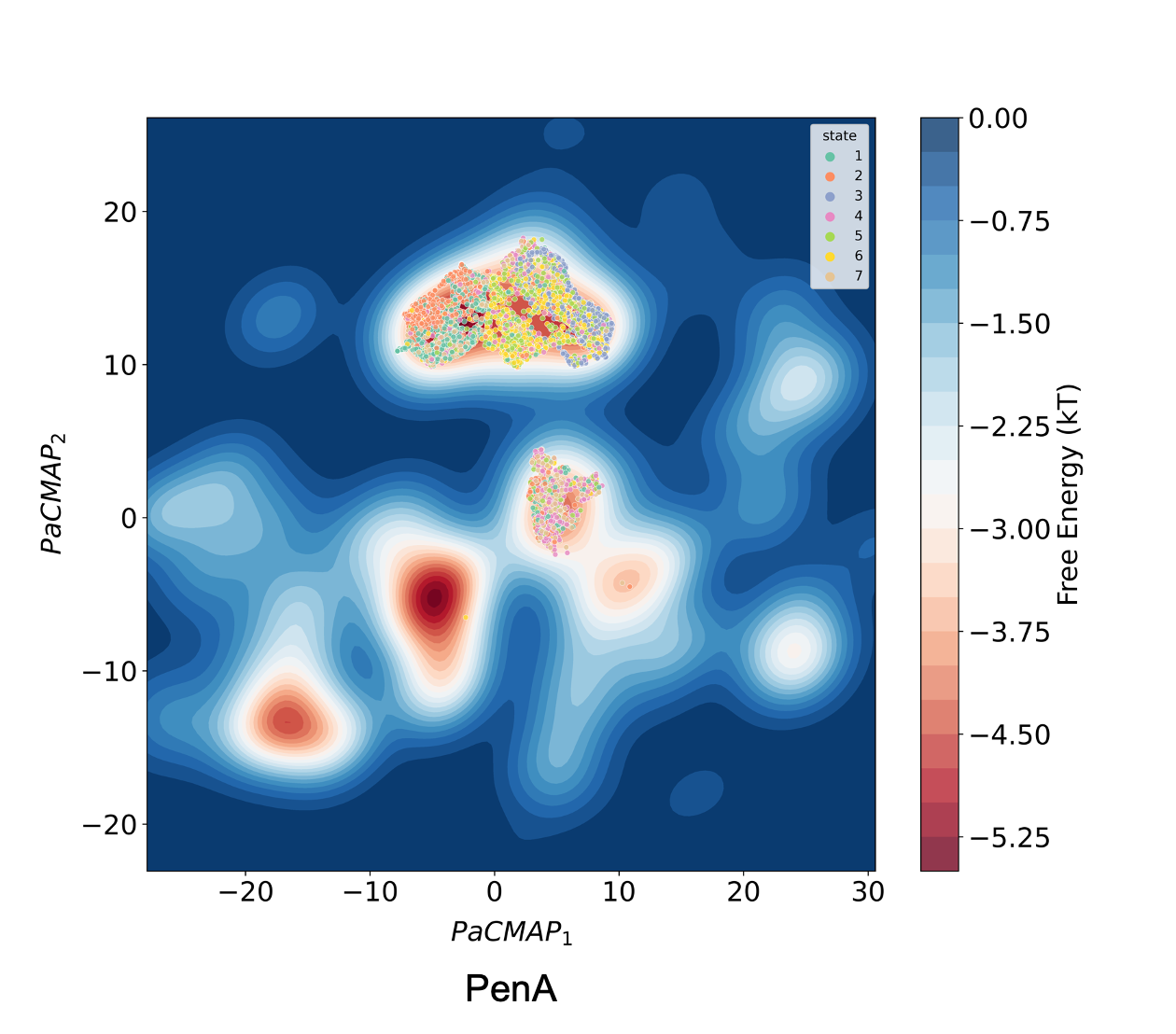

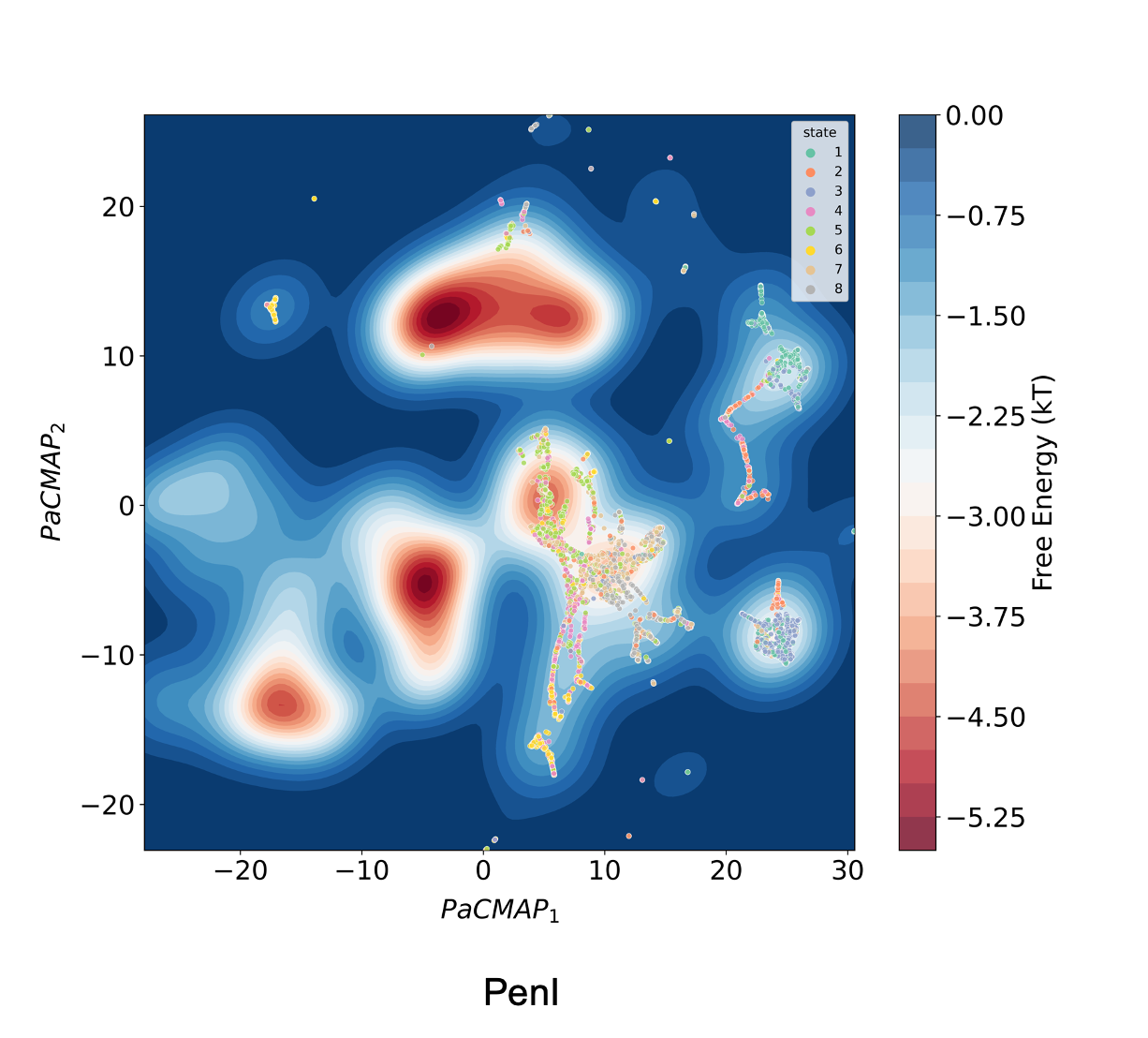

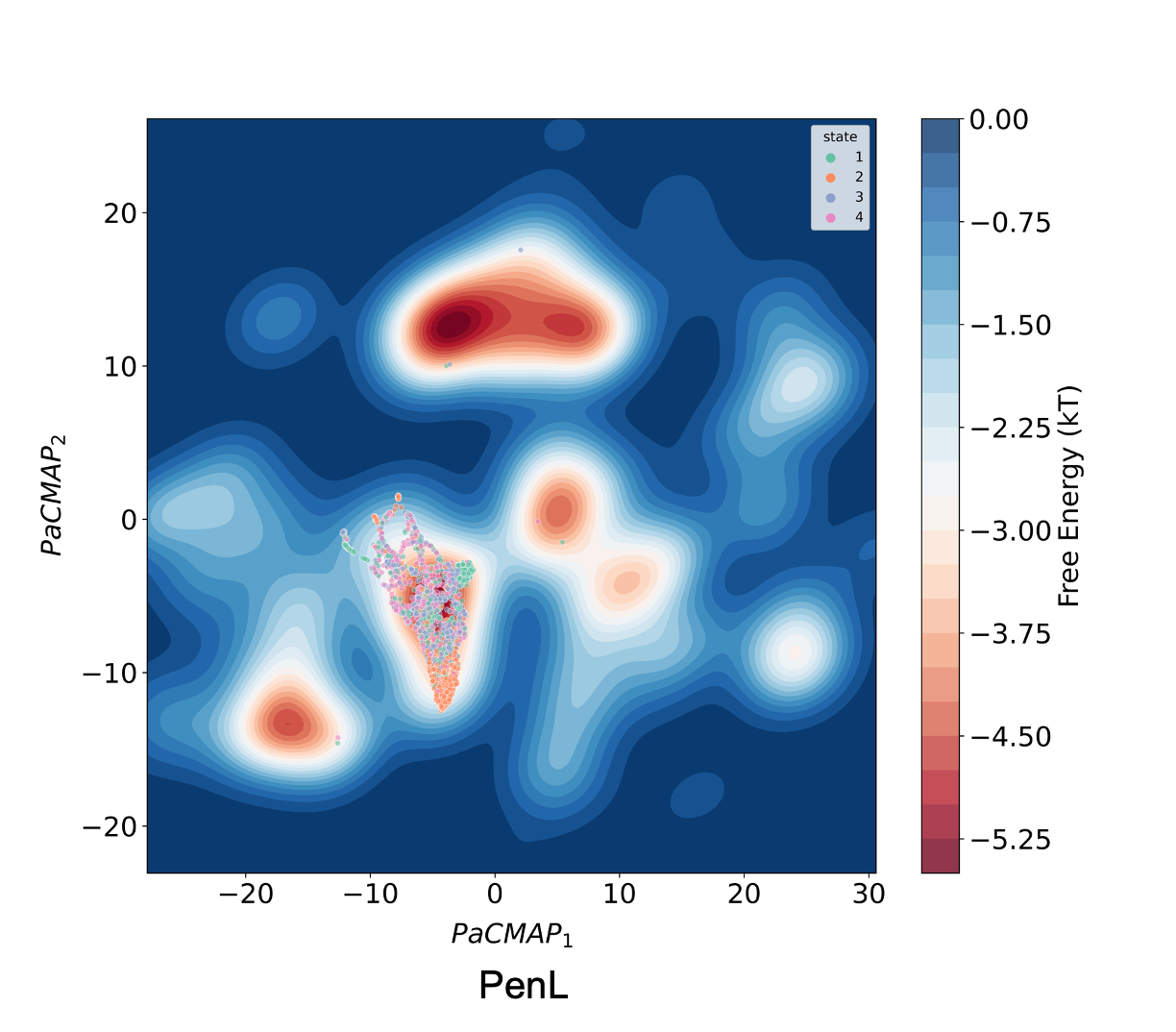

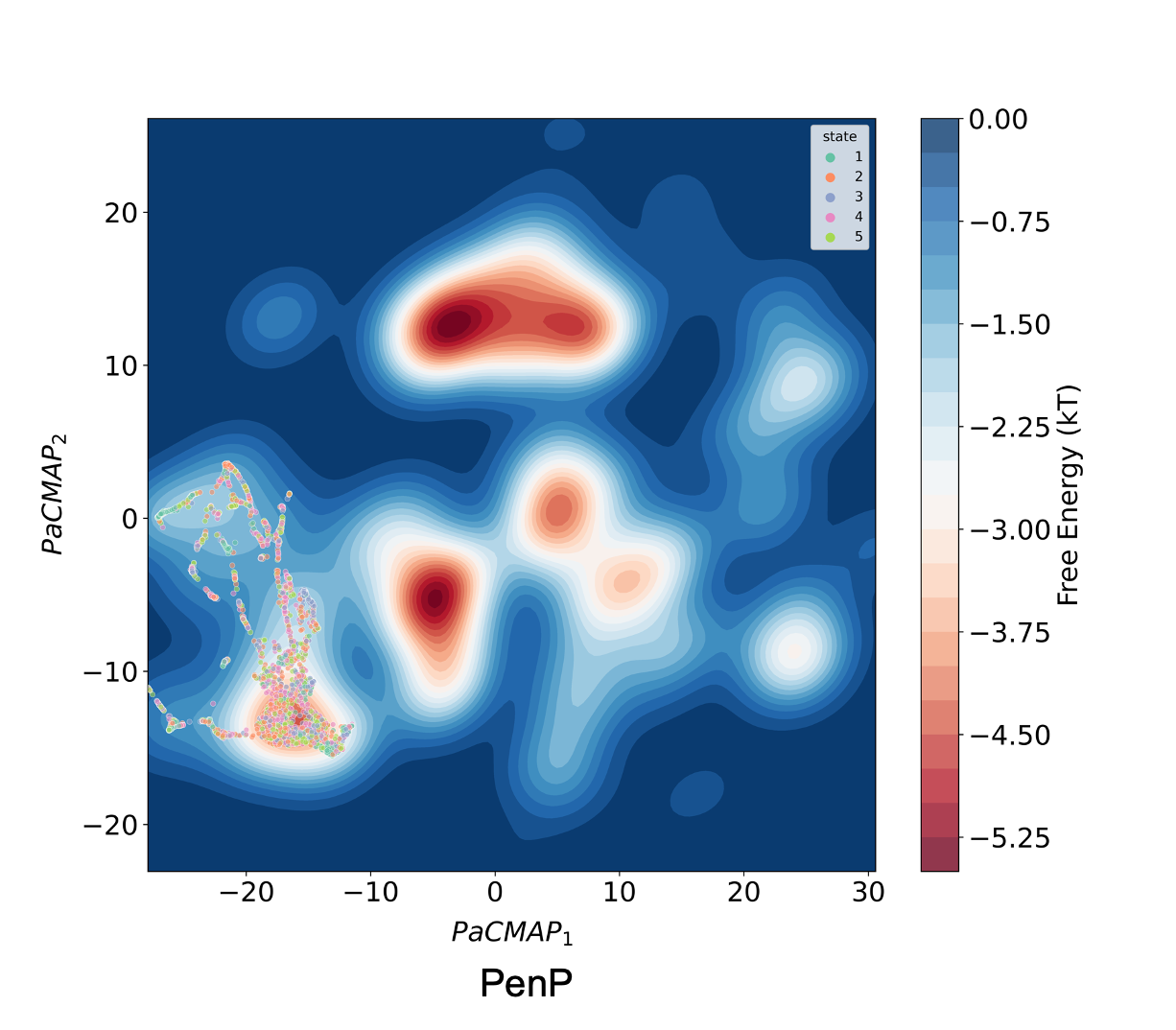

**Figure S7 Convolutional variational autoencoder (CVAE)-based deep learning analysis.**
